# Dissecting the ocular impact of SARS-CoV-2 variants: Analysis of eye globe and retina in animal models

**DOI:** 10.1101/2024.05.29.596469

**Authors:** José Igor Gomes da Silva, Victor Kersten, Livia M. Munhóz Dati, Bruna B. Lins, Tania R. Tozzeto-Mendoza, Daniela del R. F. Rodrigues, Rodrigo J. V. Barbosa, Silvia H. Lima, Lucy S. Villas Boas, Alexandre dos Santos da Silva, Khallil T. Chaim, Maria Cássia Mendes-Corrêa, Momtchilo Russo, Luciana Mirotti, Marcelo A. Pinto, Rodrigo A. P. Martins

**Author notes:** Corresponding author: **Rodrigo A. P. MARTINS**, Instituto de Ciências Biomédicas, Universidade Federal do Rio de Janeiro (UFRJ). Laboratório de Neurodesenvolvimento e Neurodegeneração, CCS, Prédio Anexo do ICB, Sala LJ.1.01, Rua César Pernetta, 1766, Cidade Universitária, 21941-902, Rio de Janeiro, RJ, Brazil.

## Abstract

Neurological and ocular manifestations were reported in COVID-19 patients and in SARS-CoV-2 infected animal models. However, the effects of SARS-CoV-2 variants of concern (VoC) on the eyes and the retina remain unclear. Here, we investigate the cellular and molecular consequences of SARS-CoV-2 VoC infection on the eye and retina in mice and hamsters. Infection with the ancestral SARS-CoV-2 and the Gamma VoC induced a subtle increase in the eye volume of K18-hAce2 mice, but no morphological alteration was observed in hamsters’ eyes. Evaluation of the ocular tropism revealed that distinct SARS-CoV-2 VoC reached the eye globe, but not the retina of K18-hAce2 mice. In contrast, SARS-CoV-2 variants are detected in the hamsters’ retina during both acute infection and after disease recovery. Despite the presence of viral RNA, no inflammation was observed in the hamster retina, as evidenced by unchanged microglial cell density and unaltered gene expression of several immune mediators. Altogether, these findings indicate a limited impact of SARS-CoV-2 variants on the eye and retina.

## Introduction

Although vaccination has substantially altered the course of the pandemic and decreased mortality rates, the (coronavirus disease) COVID-19 remains a public health issue^1^, since new variants of concern (VoC) of the severe acute respiratory syndrome coronavirus 2 (SARS-CoV-2) continue to arise and circulate^2, 3^. It is now clear that the SARS-CoV-2 infection leads to systemic diseases and a variety of neurological and ocular manifestations have been described^4–10^. Besides the initially prevalent hallmarks of olfactory and gustatory disorders, and the characteristic memory impairments of the long COVID^6, 11, 12^, ocular manifestations like conjunctivitis, epiphora and retinal lesions have been reported in COVID-19 patients^4, 13–15^. Clinical studies also reported specific impacts in the retina including microhemorrhages, lesions of ischemic pattern, optic nerve thickening and thinning of the retina^4, 14, 16–20^. While Marinho and cols reported lesions in distinct retina layers^4^, another study found retinal alterations in only one of 43 patients infected with SARS-CoV-2^21^. Therefore, a better understanding of COVID-19 ocular and retinal manifestations is necessary, since it remains unclear whether SARS-CoV-2 directly affects the retina.

The original SARS-CoV-2 (Wuhan) gave rise to Alpha, Beta and Gamma variants of concern (VoC) that were subsequently replaced by the Delta, and lately the Omicron lineages and recombinants^1^. Intranasal infection of K18-hAce2 mice with ancestral (Wuhan) SARS-CoV-2, led to the presence of viral RNA and increased content of chemokines and cytokines in the eye globe, along with retinal thickening. However, these alterations did not impair mice visual function^22, 23^. In Syrian golden hamsters, epi-ocular or intranasal infection with the ancestral SARS-CoV-2 also led to viral RNA detection in the eye and induction of efficient systemic immunity response^23, 24^. It remains to be determined whether the infection by distinct SARS-CoV-2 VoC leads to morphological and inflammatory effects in the eye or neuroinflammation in the retina.

In this work, in both the K18-hAce2 transgenic mice and in wild-type Syrian golden hamsters, we evaluated the consequences of intranasal infection with the Wuhan SARS-CoV-2 or its VoC (Gamma, Delta and Omicron/BA.1) to the eye and the retina. We show that ancestral SARS-CoV-2 and the VoC Gamma and Delta invade the eye, but not the retina in the K18-hACE2 mice. Volume measurements revealed a slight increase in the eye volume of the infected mice. In hamsters, even though viral RNA was detected in the retina, no evidence of morphological alterations was observed in the eyes. Furthermore, analysis of microglial cell density and gene expression of immune mediators (cytokines and chemokines) did not indicate retinal neuroinflammation following infection by distinct SARS-CoV-2 VoC in either model. Our findings support that infection with distinct SARS-CoV-2 VoC leads to ocular manifestations (increased eye volume), however, no evidence of retinal neuroinflammation was found.

## Materials and Methods

### Animal models and SARS-CoV-2 infection

Transgenic mice expressing the human hACE2 receptor under the control of the keratin 18 promoter (K18) (B6.Cg-Tg(K18-ACE2)2Prlmn/J)^25^ were purchased from Jackson laboratories and bred at ICTB, Fiocruz, Rio de Janeiro, Brazil. The use of K18-hACE2 mice was approved by the ethics Committee for the use of laboratory animals of ICB, USP (Ethic Protocol 4344010720, CEUA 621). Mice were kept in a specific pathogen free (SPF) Biosafety Level (BSL)-2 animal house containing environment enrichment and 2 days before virus infection were transferred to a BSL-3^26^. The use of Golden Syrian hamsters (*Mesocricetus* auratus) was approved by the ethics Committee for the use of laboratory animals of Fiocruz (CEUA LW-9/20) and was carried out in accordance with the guide for the care and use of laboratory animals of Fiocruz. Adult hamsters were provided by the breeding colony of Fiocruz, Bahia and housed in a BSL-3 animal facility (Laboratório de Experimentação Animal, Bio-Manguinhos, Fiocruz). In all experiments, the animals were conditioned in climate-controlled rooms (22°C and humidity 55 ± 5%) with a 12 h light/dark cycle with food and water *ad libitum*. Animals were treated according to animal welfare guidelines under National Legislation (law 11.794).

Transgenic mice (4 - 6 weeks-old) were intranasally infected with 0.7 × 10^5^ plaque-forming units (PFU)/ml of SARS-CoV-2 (Ancestral, Gamma or Delta) in a total volume of 30 µL (15 µL per nostril) and the infected mice were kept in BSL-3 environment for a maximum of 7 days. Euthanasia was performed in mice reaching 20% of weight loss or showing signals of suffering, such as decreased activity, piloerection, un-groomed appearance, abnormal stance with ataxia, changes in eye brightness or change in respiratory pattern. Hamsters were divided into three groups of twelve animals (three per cage). Virus infection was performed via intranasal inoculation (50 µl) with 1.0 × 10^6^ PFU/ml of the SARS-CoV-2 variants studied (Delta or Omicron) in a validated laminar flow in BSL3 facility. After the infection, three animals were randomly euthanized at 3-, 5-, 10- or 15-days post infection (dpi) and the eye was collected. Euthanasia procedures were performed by total exsanguination, with blood drained by heart puncture, under deep surgical anesthesia consisted of pre-anesthesia with xylazine (1 mg/kg) associated with ketamine hydrochloride (80 mg/kg) and tramadol hydrochloride (10 mg/kg), after pre-anesthesia they were submitted with 90 mg/kg of thiopental sodic by intraperitoneal route.

### Isolation of SARS-CoV-2 variants of concern (VoC)

The ancestral Wuhan B.1.1.28 strain (EPI_ISL_1557222, isolated from a nasopharyngeal swab taken from a patient of São Caetano do Sul, Brazil in April, 2020), the Gamma P.1 strain (EPI_ISL_1060902, obtained from a nasopharyngeal specimen of a patient from Manaus, Brazil in December, 2020) and the Delta (EPI_ISL_2938096, strain B.1.617.2, Instituto Butantan) VoC were used to infect K18-hACE2 mice as previously described^26^. For the Syrian hamster experiments, a SARS-CoV-2 Delta (AY.6) (EPI_ISL_2645417, isolated from a nasopharyngeal swab from a patient from São Luís, Brazil in May, 2021), and an Omicron (BA.1) (EPI_ISL_8430488, isolated from a nasopharyngeal swab from a patient from Jaraguá do Sul, Brazil in December, 2021) were used. Both variants were kindly provided by the Laboratório de Tecnologia Virológica (Biomanguinhos, Fiocruz) and were amplified in the Vero E6 cell line to form the inoculum.

### MRI

K18-hACE2 mice were subjected to an ex-vivo MR imaging protocol on a 7 Tesla scanner (Classic Magneton, Siemens Healthcare, Erlangen, USA) equipped with an in-house transmitting and receiving solenoid coil in the core facilities PISA of the Faculty of Medicine of the University of São Paulo (USP). A solenoid coil, designed to reduce movement during image acquisitions, housed a 50 mL falcon tube containing the paraformaldehyde-fixed mice head in 1x PBS.

Two volumetric data were acquired with 100 micrometer isotropic protocols: 3D-Turbo Spin Echo (TR/TE = 2000/122 ms; FoV = 29×59 mm; matrix = 288×576; 256 slices; slice thickness 0.10 mm; echo train length = 20; variable flip angle = 120°; number of averages = 1; acquisition time = 2h 7min) and 3D-Gradient Echo (TR/TE = 45/13 ms; FoV = 29×59 mm; matrix = 288×576; 256 slices; slice thickness 0.10 mm; flip angle = 15°; number of averages = 1; acquisition time = 56min).

Using MIPAV (https://mipav.cit.nih.gov/) software version 11.0.3, a volume of interest (VOI) for the vitreous humor region of each eye of every subject was delineated in the 3D-TSE acquisition, using the coronal plane. Subsequently, the total volume in cubic millimeters was extracted.

### Viral load quantification

For SARS-CoV-2 genome amplification an adapted protocol adapted was used^27^. In this protocol we used the forward primer ACAGGTACGTTAATAGTTAATAGCGT, reverse primer ATATTGCAGCAGTACGCACACA and probe 6FAM-ACACTAGCCATCCTTACTGCGCTTCG-QSY to detect and quantify the envelope (E) sequence of SARS-CoV-2. Thermal cycling was performed at 45 °C for 10 minutes for reverse transcription, followed by 95 °C for 10 minutes for initial denaturation and amplification with 45 cycles of 95 °C for 15 seconds and 60 °C for 45 seconds.

### RNA isolation and real-time RT-PCR

Eyes were collected, rinsed in 1x PBS, kept overnight in RNALater (Sigma Aldrich, R0901) at 4°C and stored at −80°C until processing. The full eye globe or dissected retinas were lysed in QIAzol lysis reagent (Qiagen, 79306) using a tissue homogenizer (Omni International, SKU TH115). The RNA was extracted using the RNeasy Plus Mini Kit (250) (Quiagen, 74136), following manufacturer’s instructions and quantified using a NanoDrop Lite spectrophotometer (Thermo Fisher Scientific). The cDNA was synthesized using the High-Capacity RNA-to-cDNA kit, according to the manufacturer’s instructions (Thermo Fisher Scientific, 4388950).

Real-Time RT-PCR reactions were performed in a QuantStudio 7 Flex Real-Time PCR System using SYBR green technology (PowerTrack SYBR Green Master Mix, Thermo Fisher Scientific, A46109), following the manufacturer’s instructions. Target genes and their primer sequences are listed in Table 1. Each sample was run in duplicate and only replicates varying less than 0.5 Ct between them were considered. Relative gene expression was determined applying the 2^-ΔΔCt^ method^28^, using the average β-Actin expression as reference.

**Table 1:**
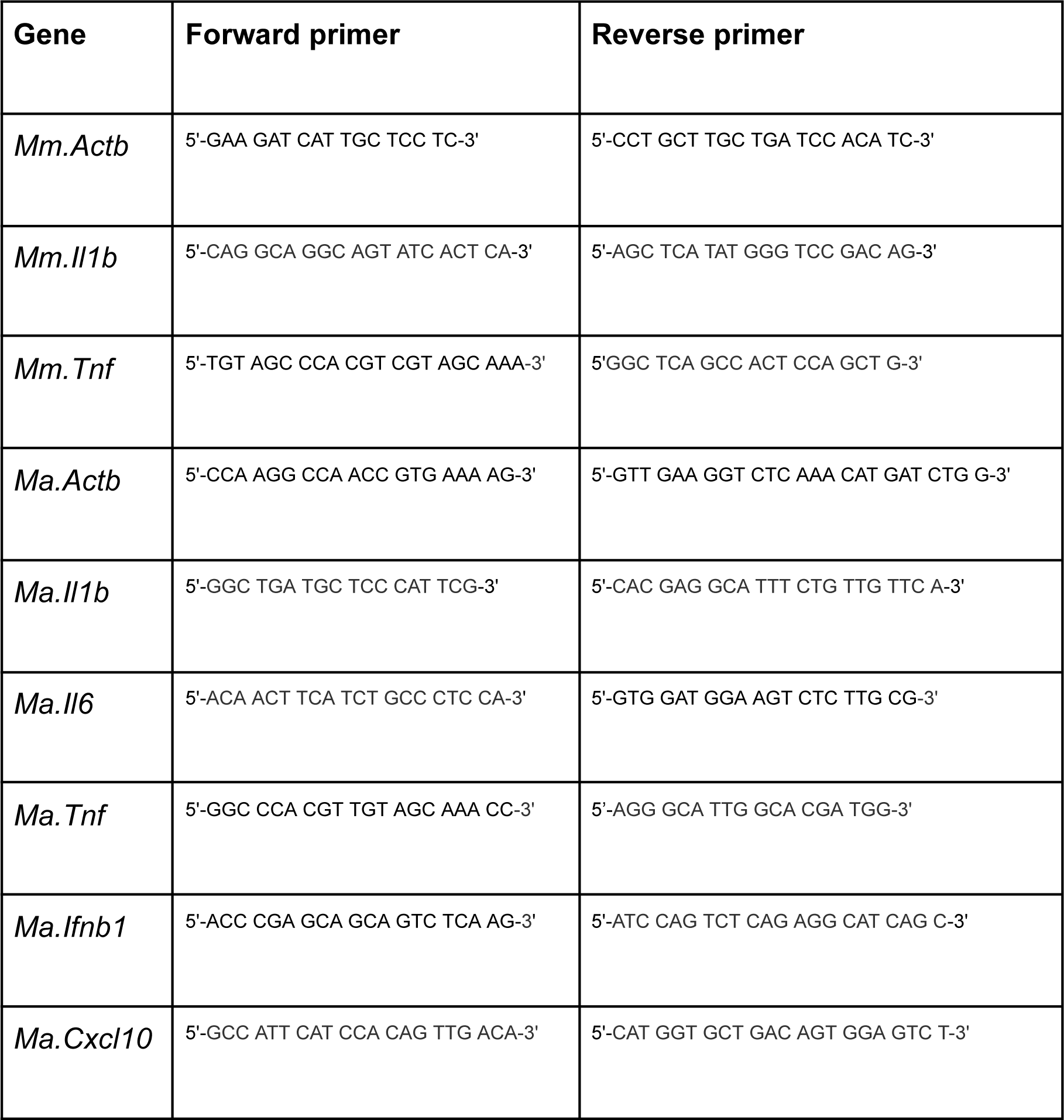
Real-time RT-PCR primer sequences.

### Eye volume measurements

The volume of Syrian hamster eyes was measured as previously described^29^. The axial length and two coronal axes (dorso-ventral and medial-lateral) of each eye were measured with a digital paquimeter and the eye volume was calculated after applying the formula (4/3× PI) × (axial length in mm) × (medial-lateral length in mm) × (dorsal-ventral length in mm).

### Histology, immunostaining, microscopy and quantification of microglia

Eyes were dissected for removal of cornea and lens and fixed in 4% paraformaldehyde (Santa Cruz Biotechnology, sc-281692) overnight, washed in 1x PBS and cryoprotected by a sucrose gradient (10%, 20% and 30% for 16h at 4°C). Then, tissues were covered on OCT (Fisher Healthcare, 4585) and frozen in liquid nitrogen. 10 µm sections were obtained in a Leica CM1850 cryostat and mounted in poly-L-lysine-coated slides (300 μg/mL, Sigma Aldrich, P2636).

Slides were washed in PBS 1X and heat-induced (microwave) antigen retrieval was performed in a 10 mM citrate buffer (pH 6.0). Slides were incubated for 1h in blocking solution [5% goat serum (Sigma Aldrich, D9023); 1% BSA (Sigma, A2153) and 0.5% Triton X-100 (Sigma, X100)]. Primary antibodies (rabbit) against Ionized calcium binding adaptor molecule 1 (Iba-1, 1:200, Wako Chemicals, 019-19741) were diluted in blocking solution and slides were incubated overnight at 4°C. Then, slides were washed in PBS 1X and incubated for 45 minutes with goat anti-Rabbit IgG secondary biotinylated antibody (1:500, Vector Laboratories, BA-1000) followed by an incubation with Cy3-conjugated streptavidin (1:300, Thermo Fisher Scientific, 434315). Nuclei were counterstained with DAPI (Sigma-Aldrich, D9542). Fluorescent images were captured using a Zeiss M2 Apotome (Zeiss Group, Germany).

For the quantification of microglial cell density, only Iba1-positive (+) cell soma that coincided with DAPI (nuclear) staining were considered. Ten retinal sections were quantified for every animal analyzed; and the length (in mm) of each section was also registered. Then, the mean of the number of Iba1+ cells per mm of retina was calculated for each animal.

### Quantification and statistical analysis

Three or more biological replicates (animals) were analyzed in each experimental group, and each dot on a plot indicates one biological replicate. Statistical significance was determined using one-way ANOVA with Dunett’s multiple comparisons test. Graph Pad Prism 9 software (La Jolla, CA, USA) was used for statistical analysis and graph generation.

## Results

### Ocular manifestations following infection by SARS-CoV-2 VoC

In a recent study, we confirmed previous findings that intranasal infection of K18-hACE2 with ancestral (Wuhan) SARS-CoV-2 leads to an acute and systemic disease characterized by weight loss, lung injury, neuroinflammation in the brain and lethality ∼7 days post infection (dpi)^23, 26, 30–32^. To investigate whether distinct SARS-CoV-2 VoC could affect eye morphology, we intranasally infected K18-hACE2 mice with the ancestral (Wuhan) or Gamma/P.1 and measured eye volume 6 dpi. Magnetic resonance imaging (MRI) and morphometric analysis revealed an increase of ∼60% in the eye volume of ancestral SARS-CoV-2- and Gamma-infected mice (Figure 1).

**Figure 1:**
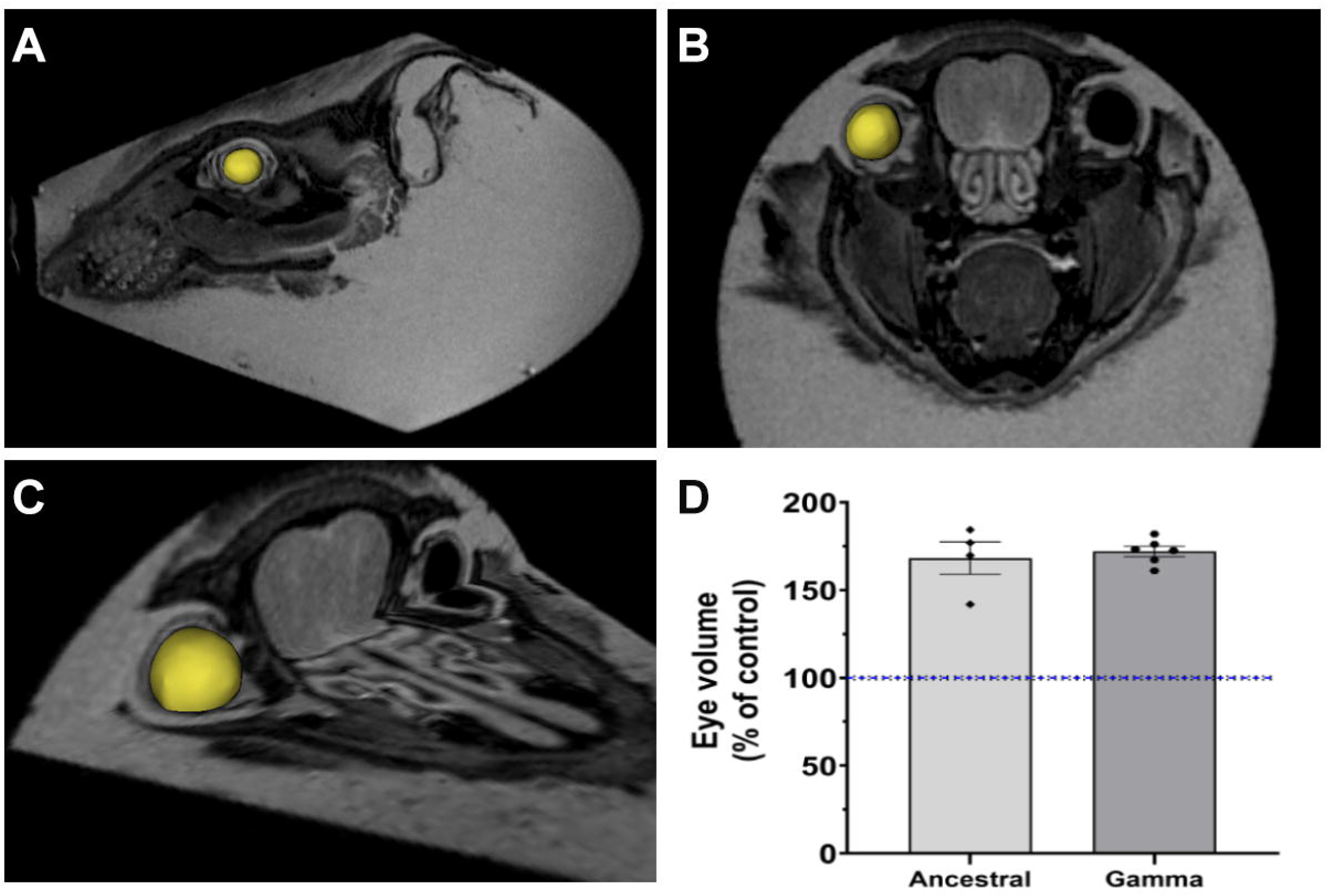
Morphometric imaging analysis revealed an increase in the eye volume of SARS-CoV-2-infected mice. (A-C) Representative images of K18-hACE2 mice head cross-sections views generated by MRI. The area occupied by the eyes in each plane (sagittal, coronal or orthogonal) is shown in yellow. (D) Eye volume measurements of control (n=10), ancestral- (n=4) and Gamma-infected (n=6) K18-hACE2 mice at 6 dpi. The blue dashed line represents the mean and the black dashed lines represent the S.E.M of non-infected (control) samples.

### Ocular tropism of SARS-CoV-2 VoC: Eye globe vs retina

ACE2 and TMPRSS2 are expressed in human ocular tissues and in other species, including mice and pigs^33, 34^. More specifically, ACE2 expression has been reported in human retinas^35, 36^. Following infection of K18-hACE2 mice with ancestral virus, SARS-CoV-2 was detected in the eye globes^23^. To analyze whether SARS-CoV-2 VoC reaches the retinal tissue, we infected K18-hACE2 with ancestral, Gamma or Delta SARS-CoV-2 and quantified viral RNA copies. We confirmed that ancestral SARS-CoV-2 spreads to the eye globes of K18-hACE2 mice. Similarly, viral RNA was detected in the eye globes following infection with both Gamma and Delta VoC. In contrast, at 4 dpi, no viral RNA was detected in K18-hACE2 retinal tissue in any of the SARS-CoV-2 VoC studied (Figure 2). These results indicate that SARS-CoV-2 reaches the eye globes, but not the retina.

**Figure 2:**
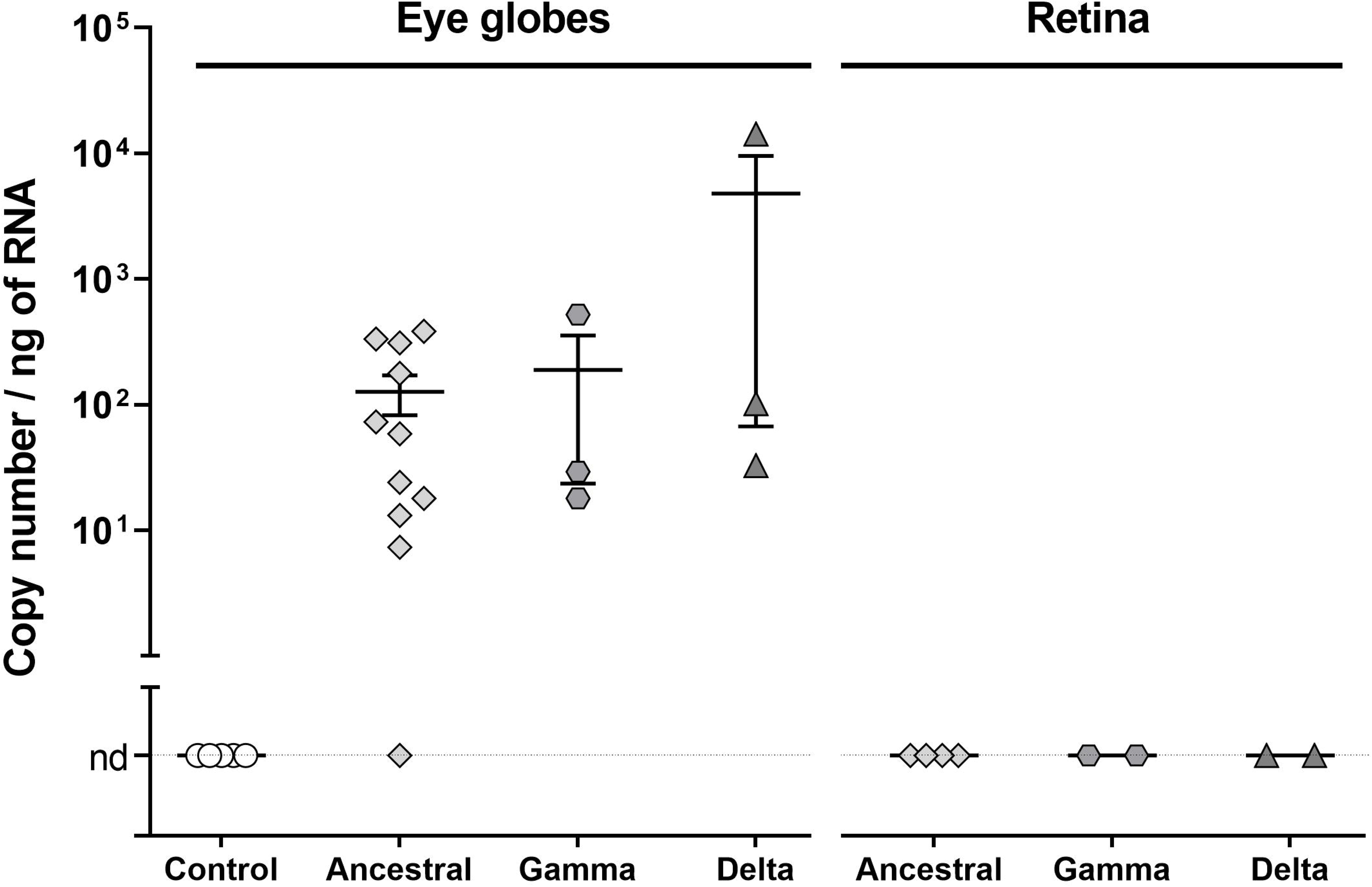
SARS-CoV-2 VoC reaches the eye globe, but not the retina of K18-hACE2 mice. RT-PCR-based quantification of SARS-CoV-2 RNA in the eye globe and in the retina of K18-hACE2-infected mice. Mice were intranasally infected with ancestral (n=4), Gamma (n=3) or Delta (n=3) SARS-CoV-2. The whole eye globe or the retina were harvested at 4dpi. The data indicate means ± S.E.M.

### Proinflammatory cytokines are not upregulated in the retina of K18-hACE2-infected mice

To determine whether SARS-CoV-2 VoC induces inflammation in ocular tissues, mRNA levels of proinflammatory cytokines were measured by real-time RT-PCR. Consistent with the viral load analysis, no alteration in the mRNA expression of *Il1b* and *Tnf* was observed in the retina or in the eye globes at 4dpi (Figure 3).

**Figure 3:**
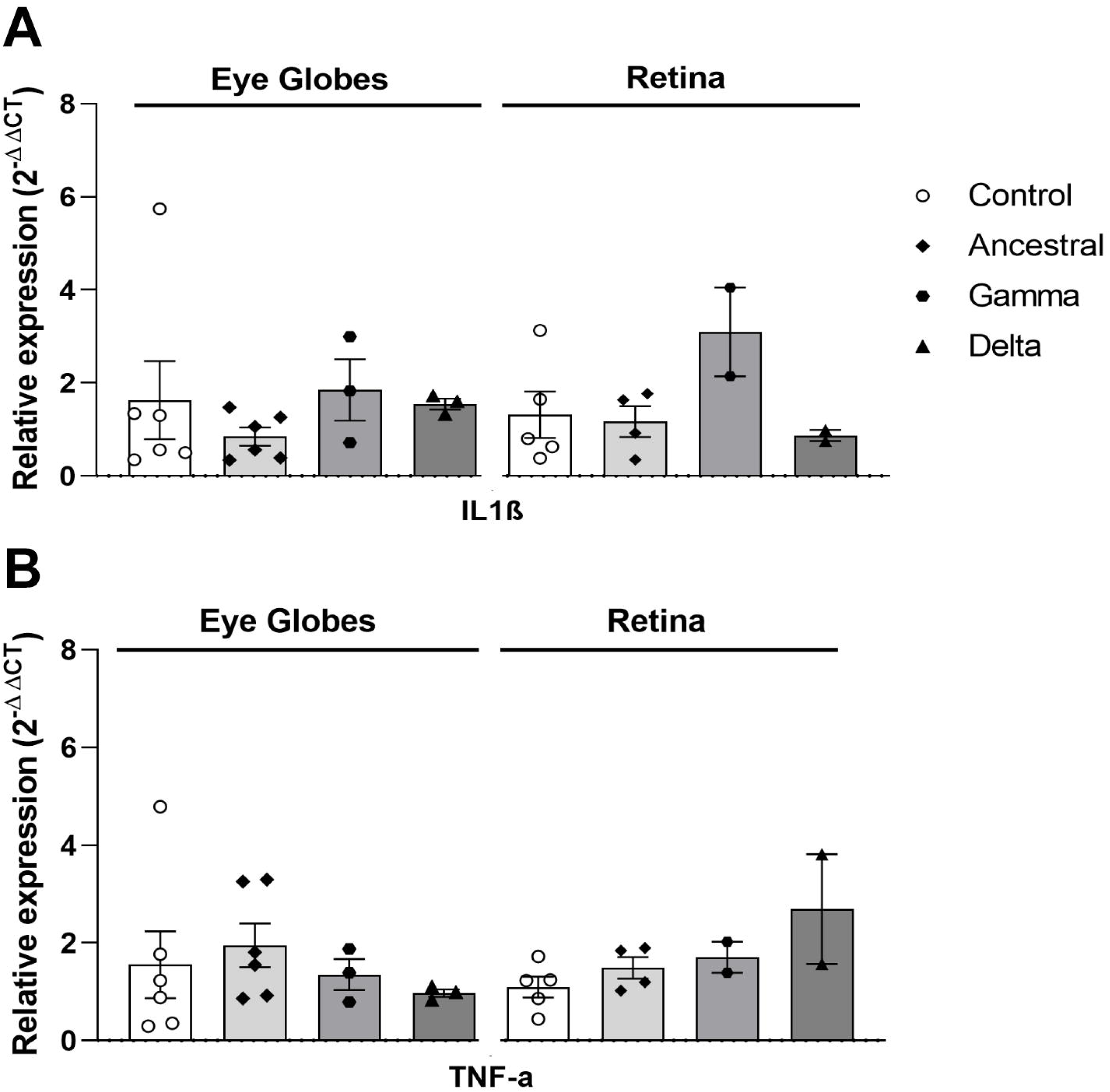
Specific SARS-CoV-2 VoC upregulates proinflammatory cytokines in the eyes, but not in the retina of K18-hAce2-infected mice. Mice were intranasally infected with ancestral (n=6), Gamma (n=3) or Delta (n=3) SARS-CoV-2. Eye globes or retina were harvested at 3 and 5 dpi. Relative expression of *Il1b* and *Tnf* mRNAs in the eye globes and in the retina of K18-hACE2-infected mice at 4dpi are shown. The data indicate means ± S.E.M.

### Lack of ocular manifestations or retinal inflammation in hamsters infected with SARS-CoV-2 variants

hAce2 overexpression in K18-hAce2 mice (8 transgene copies)^25^ raised concerns that the observed neuropathology might not reflect human COVID-19. Syrian golden hamsters are also an excellent model to study COVID-19 due to a consistent pathogenesis, efficient immune response, and low mortality. Several studies showed that intranasal infection of hamsters with the ancestral SARS-CoV-2 leads to a transient body weight loss (∼10%) ^23, 37–39^. To analyze whether SARS-CoV-2 VoC induce a similar disease course, we intranasally infected hamsters with Delta or Omicron VoC. A comparable body weight loss at 6-8 dpi as well as a recovery to initial weight around 14 dpi were observed (Figure 4A). To evaluate whether SARS-COV-2 VoC could impact eye morphology in hamsters, we measured eye volume during acute COVID-19 and following recovery (3, 5, 10 and 15 dpi). No alteration in eye volume was induced by any of the SARS-COV-2 VoC studied here, indicating that COVID-19 does not induce macroscopic ocular manifestations in hamsters (Figure 4B).

**Figure 4:**
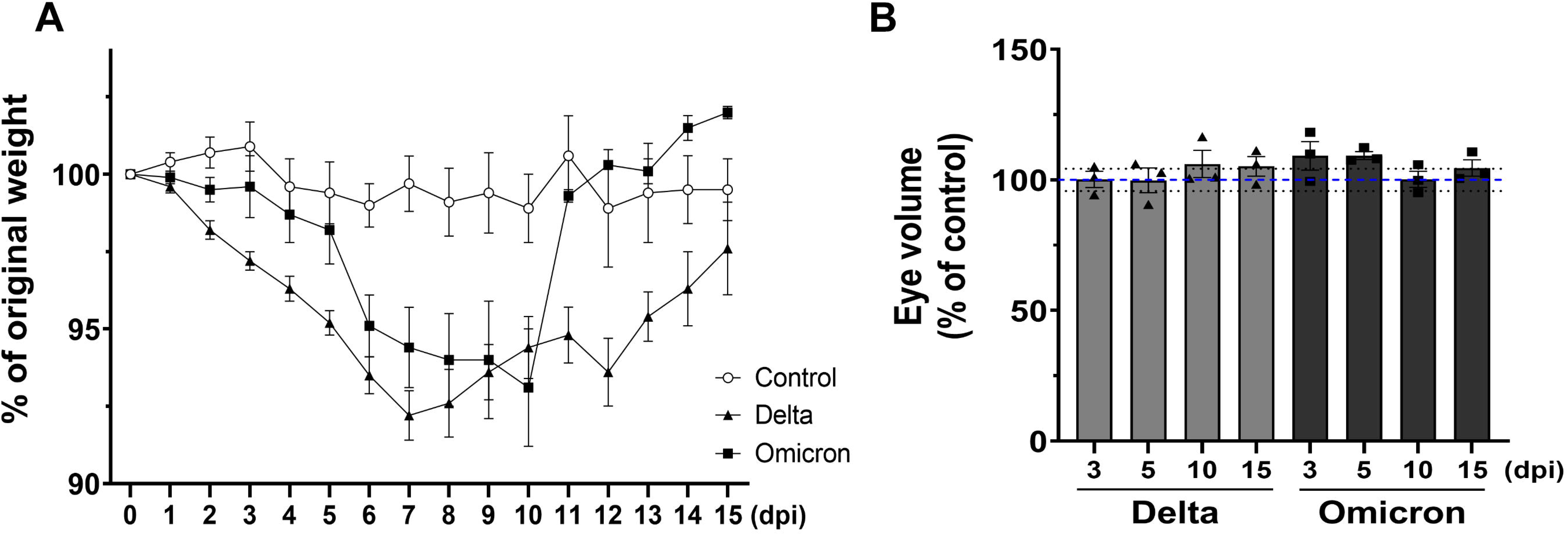
SARS-CoV-2 variants do not affect the eye volume of the Syrian golden hamsters. Syrian hamsters were intranasally infected with 10^5^ PFU of Delta (n=3) or Omicron (n=3) SARS-CoV-2 VoC. (A) Body weight measurements are shown as a percentage of the initial body weight at the indicated days post-infection (dpi). (B) Eye volume measurements in hamsters infected with 10^5^ PFU of Delta- (n=3) or Omicron (n=3). Eyes were harvested and measured 3, 5, 10 or 15 dpi. The blue dashed line represents the mean and the black dashed lines represent the S.E.M of non-infected (control) samples. The bars indicate the means ± S.E.M of infected hamsters.

In hamsters, SARS-CoV-2 RNA was detected in the eye globes following infection with the ancestral virus^23^. Moreover, different SARS-CoV-2 VoCs were shown to invade distinct regions of hamsters CNS^8^. To analyze whether SARS-CoV-2 VoC reach the retina, following intranasal infection with Delta or Omicron VoC, we determined the copy number of SARS-CoV-2 RNA in the retina (3, 5, 10 and 15 dpi). Delta and Omicron SARS-CoV-2 RNA were detected in ∼75% or 60% of the hamster retinas analyzed, respectively. Consistent with previous CNS studies^8^, viral RNA was detected in the retina during acute disease (5dpi) and following recovery (10 and 15 dpi) for both Delta and Omicron VOC (Figure 5). These data indicate that distinct SARS-CoV-2 VoC reaches hamsters’ retina.

**Figure 5:**
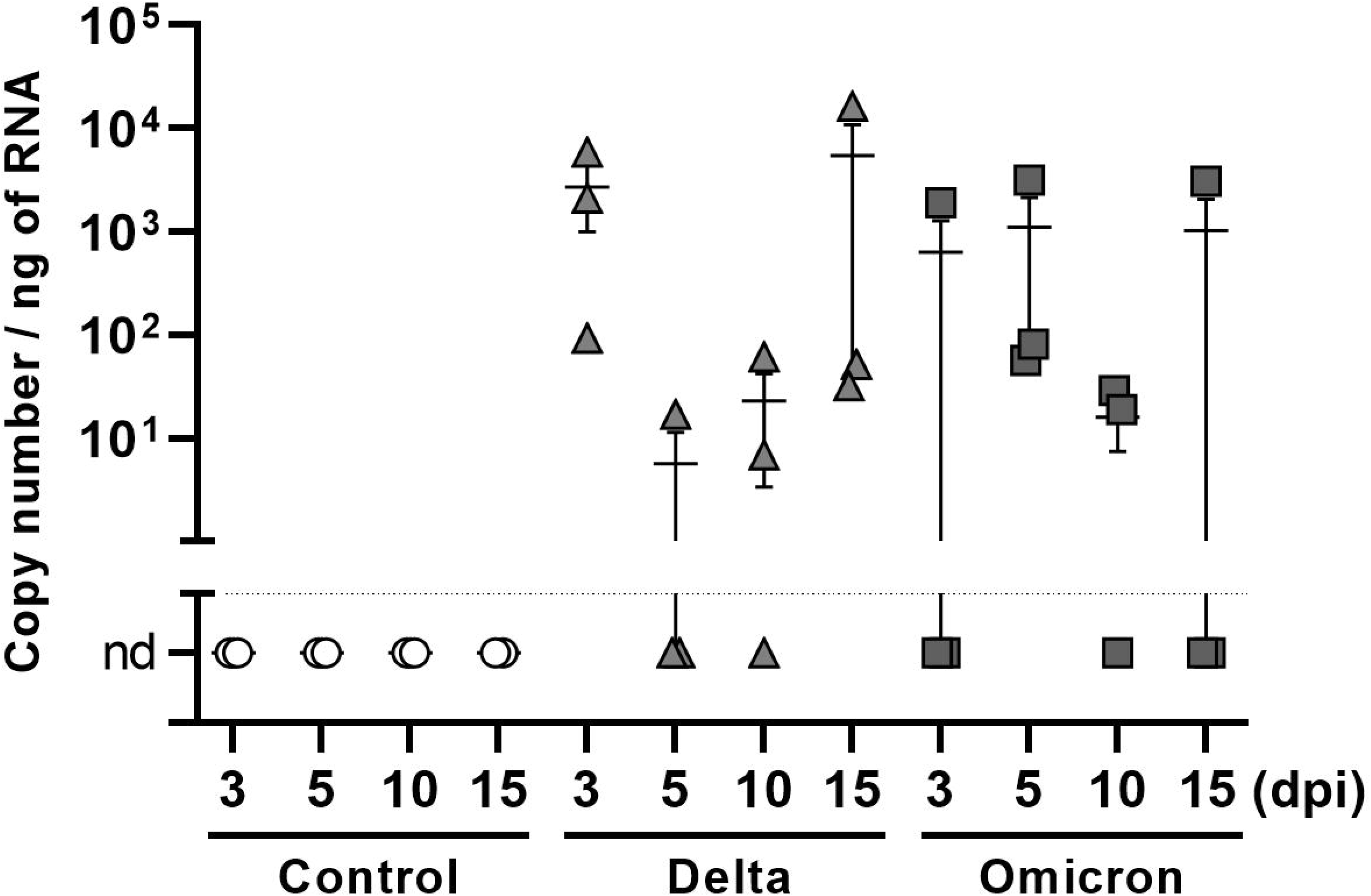
Detection of SARS-CoV-2 variants RNA in the retina of Syrian golden hamsters. Real-time PCR-based quantification of SARS-CoV-2 RNA (gene E) in the retina. Syrian hamsters were intranasally infected with 10^5^ PFU of Delta (n=3) or Omicron (n=3) SARS-CoV-2 VoC and retinas were harvested 3, 5, 10 or 15 dpi. Viral load quantification indicates the number of SARS-CoV-2 RNA copies per ng of RNA. The data indicate means ± S.E.M.

Microglia is the main innate sensor of viral infections in the CNS^40^ and SARS-CoV-2 infection induces microglial cells activation and proliferation in the CNS^41–43^. To assess possible inflammatory consequences of Delta or Omicron VoC infection within the retina, we investigated whether SARS-CoV-2 infection could affect retinal microglia. For this, we harvested retinas during the acute phase of the infection (5 dpi), stained for the microglial marker Iba1 and quantified the density of Iba1+ cells. Intranasal infection with SARS-CoV-2 Delta or Omicron VoC does not impact microglial cells density in the hamster retina. This finding indicates no proliferation or recruitment of microglia occurred suggesting the lack of neuroinflammation in the retina (Figure 6).

**Figure 6:**
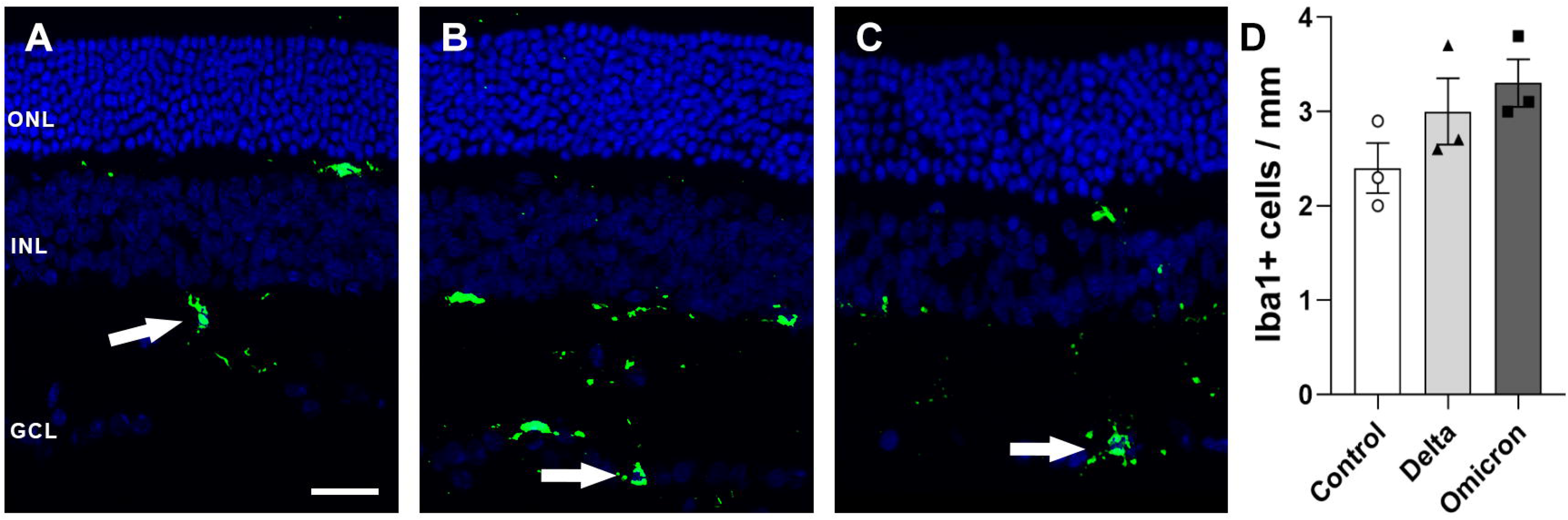
Intranasal infection with SARS-CoV-2 variants does not affect microglial cells density in the retina of Syrian golden hamsters. (A-C) Representative confocal images of Iba1-positive (+) cells (green) in the retina. Arrows point to Iba1 + cells in each group. Syrian hamsters were intranasally infected with 10^5^ PFU of Delta (n=3) or Omicron (n=3) SARS-CoV-2 VoC and retinas were harvested at 5 dpi. (D) Quantification of Iba1 + cells density in the retina of control, Delta- or Omicron-infected hamsters (n=3) 5 dpi. The data indicate means ± S.E.M.

Different SARS-CoV-2 VoC cause neuroinflammation and deregulate immune mediator’s gene expression in the olfactory bulb and other brain areas in hamsters^44, 45^. To further explore the impact of SARS-CoV-2 VoC and evaluate possible inflammatory response in the retina, we examined chemokines and cytokines mRNA expression by real-time RT-PCR. Consistent with the lack of microglial activation, no alteration in the expression of *Il1b, Il6, Tnf, Ifnb1* and *Cxcl10* was observed following infection by the Delta or Omicron SARS-COV-2 VoC (Figure 7). Therefore, despite the detection of viral RNA SARS-CoV-2 variants do not impact eye morphology or induce retinal inflammation in hamsters.

**Figure 7:**
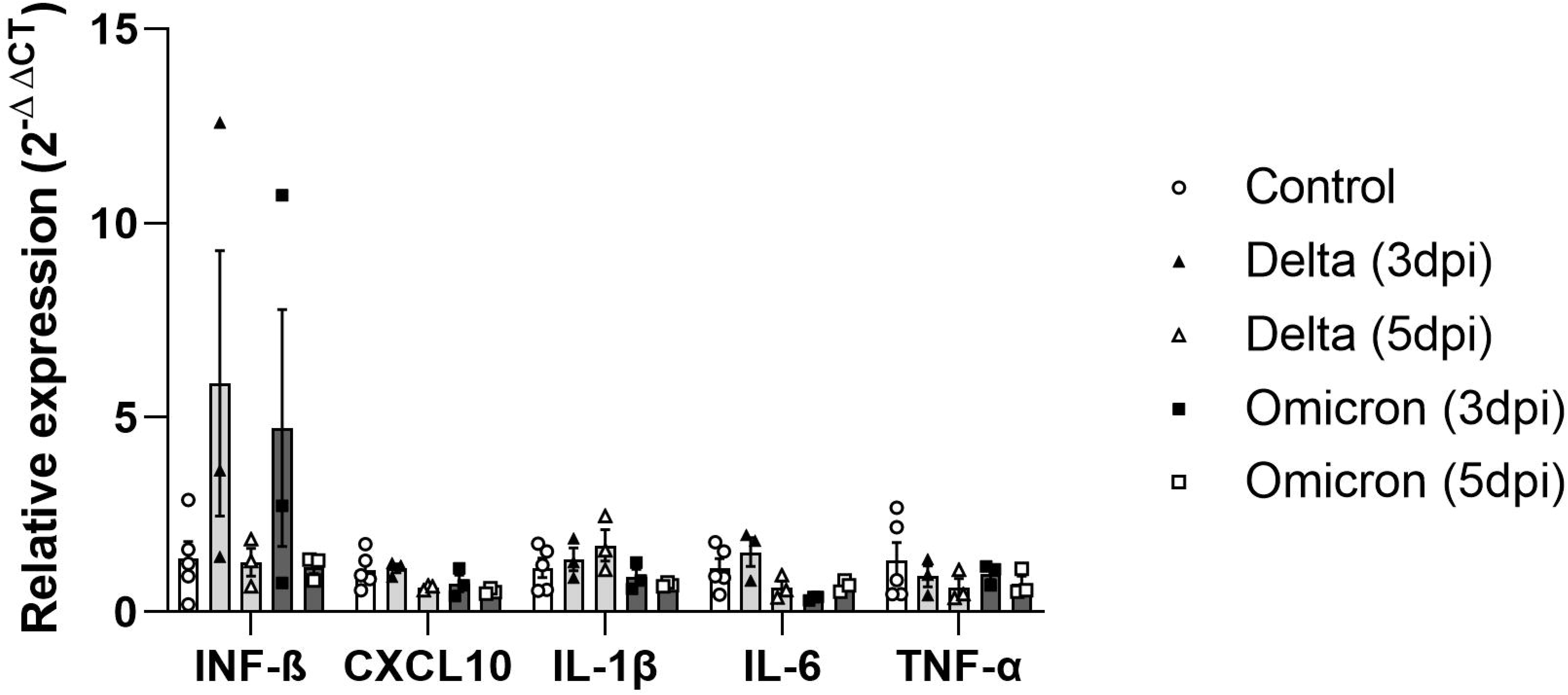
Infection with SARS-CoV-2 variants does not affect cytokines and chemokines expression in the retina of Syrian golden hamsters. Syrian hamsters were intranasally infected with 10^5^ PFU of Delta (n=3) or Omicron (n=3) SARS-CoV-2 VoC. Retinas were harvested at 3 or 5 dpi. Expression of *Il1b, Il6, Tnf, Ifnb1 and Cxcl10* mRNAs was not altered in retinal tissue of control, Delta- or Omicron-infected hamsters at 3 and 5 dpi. The data indicate means ± S.E.M.

## Discussion

Although it is clear that COVID-19 affects the brain and other nervous system organs in humans and in animal models^5–7, 10^, the cellular and the molecular consequences of SARS-CoV-2 infection to the eyes and to the retina remains unclear. Few studies published data about the effects of the original SARS-CoV-2 (Wuhan) to the eyes^23^; however, to our knowledge, no information concerning the impact of SARS-CoV-2 VoC infection on the eye globes and retina is available. Overall, our results show that SARS-CoV-2 VoC differentially reach ocular tissues and induce subtle morphological effects in the eyes of infected K18-hAce2 mice. Notably, analysis of microglia reactivity and proinflammatory cytokine expression in both animal models did not indicate SARS-CoV-2 VoC-induced retinal inflammation.

Ocular manifestations induced by viral infections have been previously reported^46^. Influenza A infection upregulates inflammation-associated genes in human retinal microvascular endothelial cells^47^. Zika virus infection induces severe chorioretinal atrophy of the macula and pigmentary changes^48^. In humans, SARS-CoV-2 infection led to choroid and retinal thickening along with acute retinal necrosis^18, 19, 23^. We demonstrate that infection by SARS-CoV-2 VoC induces subtle morphological alterations in the eyes of K18-hACE2 mice, while no difference in the eye volume was observed following infection of Syrian golden hamsters.

The ACE2 receptor and TMPRSS2 are expressed in different ocular tissues, including the cornea, conjunctiva and retina in humans and other species^33–35^. Consistently, SARS-CoV-2 ocular tropism has been described in humans and animal models^23, 24^. SARS-CoV-2 nucleocapsid protein was detected in the eye globe of a COVID-19 patient that exhibited ocular manifestations^49^. Following intranasal infection with the ancestral virus, SARS-CoV-2 was found in the eye globes of both K18-hACE2 mice and golden hamsters^22, 23^. Further, SARS-CoV-2 RNA and spike protein were also specifically detected in the retina of K18-hACE2 mice^23, 30^. Here, we found SARS-CoV-2 RNA of all the VoCs studied in hamsters’ retina. In contrast, viral load assessment showed that ancestral, Gamma and Delta SARS-CoV-2 VoC reach the eye, but not in the retina of K18-hACE2 mice. Therefore, despite the different pattern across the studied species, in Syrian golden hamsters both the retinal tropism and the body weight loss is conserved among the distinct SARS-CoV-2 VoC.

In contrast to our findings in mice, SARS-CoV-2 RNA was detected in the retina of infected hamsters. Considering infections of either Delta or Omicron VoC, viral mRNA was detected in 75% recovered animals (10 and 15 dpi). Similar observations were previously reported in other CNS regions (olfactory bulb and cerebral cortex)^8^. In distinct brain regions, regardless of the VoC considered, neuroinflammation is a common feature as revealed by the upregulation of several cytokines and chemokines^8, 44^. In the hamster retina, despite the presence of viral RNA, microglial cells density and the gene expression of several immune mediators remained unaltered, supporting lack of retinal inflammation in hamsters infected with distinct SARS-CoV-2 variants. This could be explained by the differences of viral load between brain and retina, or SARS-CoV-2-induced neuroinflammatory response may be differently suppressed in these CNS regions. Moreover, even though brain and retina neuroinflammation profile is different, SARS-CoV-2 RNA remains detectable in both regions after disease recovery. Overall, our findings reveal a limited impact of SARS-CoV-2 variants to the eye and in the retina and follow-up studies will be important to better understand ocular abnormalities in acute and long COVID-19.

## Declarations

### Ethics approval and consent to participate

All authors declare consent to publish.

### Availability of data and materials

All authors agree to make data and materials available.

### Competing interests

The authors declare that no competing interests exist.

### Funding

This work was supported by grants from The International Retinal Research Foundation (IRRF 2021 to R.A.P.M.); Fundação de Amparo à Pesquisa do Estado do Rio de Janeiro (FAPERJ) (E-26/211.711/2021 to R.A.P.M.); #HCComVida initiative of the HCFMUSP to K.C.; FAPESP (2021/12502-7 to M.R.); Programa Inova Fiocruz (VPPCB-005-FIO-20-2-54 to L.M.), parliamentary budget from Dep. Davi Miranda (39540021 to L.M.) and a CAPES fellowship (88887.919697/2023-00 to JIGS).

### Author contributions

JIGS: performed research, analyzed data

VK: performed research

LMMD: performed research

BBL: performed research

TRT: performed research, analyzed data

DRFR: performed research

RJVB: performed research, analyzed data

SHL: performed research

LSVB: performed research

ASS: performed research, analyzed data

KTC: performed research, analyzed data, funding

MCM: performed research

MR: performed research, funding

LM: provided K18-hACE2 mice, funding

MAP: performed research

RAPM: performed research, designed research, analyzed data, wrote the paper, funding

## Acknowledgements

We thank Gildo de Brito Souza for technical assistance, Flavia Tatiana Guimarães Fraga for administrative assistance and Dr. Grasiella Ventura Matioszek for assistance in the confocal microscopy facility of the Instituto de Ciências Biomédicas (ICB, UFRJ).

## Notes

### Competing Interest Statement

The authors have declared no competing interest.

